# Timing of methamphetamine exposure during adolescence differentially influences parvalbumin and perineuronal net immunoreactivity in the medial prefrontal cortex of female, but not male, rats

**DOI:** 10.1101/2023.08.25.554911

**Authors:** Amara S. Brinks, Lauren K. Carrica, Dominic J. Tagler, Joshua M. Gulley, Janice M. Juraska

## Abstract

Adolescence involves significant reorganization within the medial prefrontal cortex (mPFC), including modifications to inhibitory neurotransmission mediated through parvalbumin (PV) interneurons and their surrounding perineuronal nets (PNNs). These developmental changes, which result in increased PV neuron activity in adulthood, may be disrupted by drug use resulting in lasting changes in mPFC function and behavior. Methamphetamine (METH), which is a readily available drug used by some adolescents, increases PV neuron activity and could influence the activity-dependent maturational process of these neurons. In the present study, we used male and female Sprague Dawley rats to test the hypothesis that METH exposure influences PV and PNN expression in a sex- and age-specific manner. Rats were injected daily with saline or 3.0 mg/kg METH from early adolescence (EA; 30-38 days old), late adolescence (LA; 40-48 days old), or young adulthood (60-68 days old). One day following exposure, effects of METH on PV cell and PNN expression were assessed using immunofluorescent labeling within the mPFC. METH exposure did not alter male PV neurons or PNNs. Females exposed in early adolescence or adulthood had more PV expressing neurons while those exposed in later adolescence had fewer, suggesting distinct windows of vulnerability to changes induced by METH exposure. In addition, females exposed to METH had more PNNs and more intense PV neuron staining, further suggesting that METH exposure in adolescence uniquely influences development of inhibitory circuits in the female mPFC. This study indicates that the timing of METH exposure, even within adolescence, influences its neural effects in females.

## 1. Introduction

Adolescence is the final period of neurodevelopment, marked by significant reorganization within prefrontal cortical gray matter in humans [1]. Adolescence is also a period of enhanced neural vulnerability to environmental perturbations and is associated with increased onset of psychiatric illness and illicit substance use [reviewed in 2]. From rodent models, it is known that reorganization within the medial prefrontal cortex (mPFC) during adolescence includes modifications to inhibitory neurotransmission mediated through parvalbumin (PV) inhibitory interneurons. These extensive modifications result in increased PV neuron output and changes to the excitatory/inhibitory balance [3,4]. In adults, commonly abused psychostimulant drugs (cocaine, methamphetamine) have been shown to increase PV neuron activity, intrinsic excitability, and alter PV expression [5–7]. Interestingly, adolescent, but not adult, exposure to cocaine decreases GABAergic inhibition and blocks the developmental increase in PV expression [8]. How well this extends to other psychostimulants, such as methamphetamine (METH) is not known nor is it clear if this would apply to females, since all the above studies were done in males.

Work from our laboratory and others have shown age-related gains in perineuronal nets (PNNs), which may contribute to increased PV cell output [9,10]. PNNs are extracellular aggregations of matrix proteins that increase PV cell firing rate and maturation, making them a potential regulator of mPFC development and plasticity [11,12]. Interestingly, our laboratory discovered a transient decrease in PNN expression in the female mPFC following pubertal onset [10]. Given PNNs limit plasticity, this transient decrease could result in post-pubertal vulnerability of the mPFC in females specifically. Adult exposure to cocaine has been shown to alter PNNs [6] and modulate the effects of PNN removal on the intensity of PV [13]. The effects of METH on PNN expression have yet to be explored, and additionally there is a lack of understanding about the potential influence of METH on female GABAergic neurotransmission.

METH is a longer acting stimulant than cocaine [14] and it could potentially have greater effects on the adolescent cortex. Potential effects of METH use during adolescence are especially important because use of METH during human adolescence has been associated with increased long-term negative consequences, including increased problematic use, higher relapse rates, and a quicker transition to uncontrolled use compared to adults [15,16]. Likewise, during self-administration adolescent rats escalate METH intake to a greater extent than adults [17–19]. Females are particularly vulnerable, with adolescent females escalating METH intake to a greater [18] or equal extent [19] compared to male adolescents. Additionally, impaired cognitive flexibility in adult females, but not males, has been found after METH self-administration during adolescence [20]. This all suggests that adolescence may be a time of increased negative consequences of stimulant use, particularly for females, which coincides with the development of PV neurons and PNNs in the mPFC. Timing of exposure within adolescence may also be important since many structural changes occur around female puberty [10, 21], and both sexes are behaviorally more sensitive to stress peri-pubertally in adolescence [22].

To assess the influence of METH on PV interneurons and PNNs in the mPFC, we exposed male and female Sprague Dawley rats to METH during either early or late adolescence, or in adulthood. We chose noncontingent drug exposure rather than self-administration so that we had better control over the timing and dose of drug exposure and could target specific times around female and male puberty during adolescent development. We hypothesized that METH exposure in adulthood would increase PV and PNN expression in the mPFC similar to previous literature. However, we also expected METH exposure in adolescence to disrupt the PV neuron and PNN developmental trajectory, potentially resulting in decreased PV and PNN expression. We also hypothesized that the magnitude of effects observed will be larger in females and animals exposed around peri-pubertal onset.

## 2. Methods

### 2.1 Animals and Housing

Sprague-Dawley rats (obtained from Envigo) were bred in the Psychology vivarium and 12 litters, across four breeding cohorts, were used. The day of birth was marked as postnatal day (P) 0. At weaning (P22), male and female pups were assigned to an exposure group counterbalanced across litter. Rats were double or triple housed in a temperature-controlled room with a reverse 12:12 light/dark cycle (lights off at 0900). Food and water were available *ad libitum*. Rats were weighed daily at 0800 starting at P25 to acclimate them to handling and to minimize injection stress. Daily checks for puberty started at P30 and ended when puberty was reached. For females, vaginal opening was used since it marks when levels of luteinizing hormone and ovarian hormones increase denoting female rat puberty [23]. In males, separation of the prepuce from the glans penis was used as an indicator, given this process is dependent on the high levels of androgens released at male puberty [24]. All animal procedures were performed in accordance with guidelines of the National Institutes of Health and the university’s animal care committee’s regulations.

### 2.2 METH Exposure and Tissue Collection

Rats were given once daily injections of 3.0 mg/kg methamphetamine HCl (Research Triangle Institute; distributed by The National Institute on Drug Abuse) or saline via an i.p. injection for 9 days in either early adolescence (EA), P30-38, late adolescence (LA), P40-48, or adulthood, P60-68 (Fig. 1). Injections occurred within the first hour of the dark cycle (0900-1000).

Between 24 and 28 h following the last injection (0900-1300; lights off at 0900), rats were deeply anesthetized using sodium pentobarbital i.p., followed by a cardiac perfusion using 0.1M phosphate buffered saline (PBS) and 4% paraformaldehyde as a fixative. Brains were post-fixed in 4% paraformaldehyde for 24 h and then submerged in a 30% sucrose solution for 72 h at 4 C. The mPFC was then coronally sectioned using a freezing microtome and stored at -20 C in a cryoprotectant solution of 30% glycerol, 30% ethylene glycol, 30% deionized water, and 10% 0.2M PBS. Tissue was coded with random IDs to ensure experimenters were blind to age, sex, and treatment for all tissue processing and analysis.

**Fig. 1:**
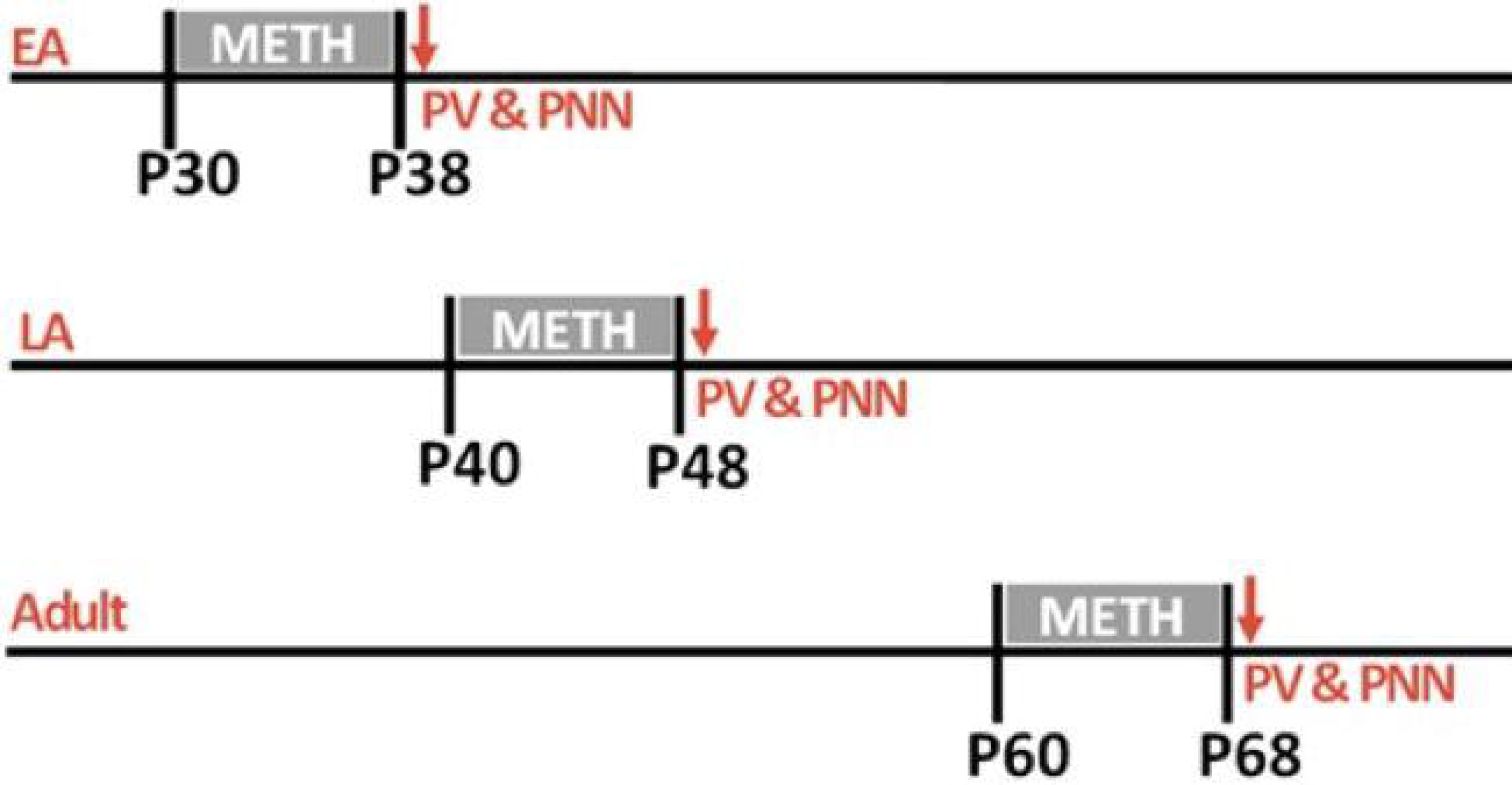
Study timeline. P = postnatal day; EA = early adolescence; LA = late adolescence. Tissue was harvested 24 h following final METH exposure to quantify PV & PNN expression.

### 2.3 Volume Quantification

To compute volume of the mPFC, every fourth section was mounted on charged microscope slides and allowed to dry before Nissl staining. Next, tissue was placed in increasing concentrations of ethanol followed by 0.2M PBS to re-hydrate the tissue. Slides were then placed in 1% periodic acid followed by Methylene Blue/Azure II, rinsed with 0.2M PBS and dehydrated using increasing concentrations of ethanol, placed in Citrisolv and coverslipped using Permount.

Volume was assessed using StereoInvestigator under a 2.5x objective, resulting in 25x magnification. The area of the mPFC was traced on each section of tissue as in our previous work [25,26]. The anterior boundary of the mPFC was denoted when the white matter of the forceps minor was evident. The most posterior section traced was the section before the appearance of the genu of the corpus callosum. The dorsal boundary of the mPFC was marked by thinning of layer 1 along with cell orientation. The ventral boundary of the mPFC was denoted by loss of visible cortical layers. The mPFC was further subdivided into its subregions, the prelimbic (PL) and infralimbic (IL) cortices. This boundary is evident by loss of gradation between layer II and III. To obtain the z-depth of the mPFC, the average thickness was sampled from 15 sites per animal using StereoInvestigator under 630x magnification and oil immersion. From these parcellated boundaries, the area of each sampled slice was calculated, multiplied by the average section thickness multiplied by 4, and then all volumes of sampled tissue were summed to obtain the entire PL and IL volume for each animal.

### 2.4 Immunofluorescent Labeling and Imaging

For each subject, three sections containing mPFC (from bregma +3.00 to +2.50) were stained for PV and PNNs. Bregma was controlled as PV may vary across the anterior-posterior gradient. First, sections were washed in tris buffered saline (TBS) in triplicate followed by a 1-h incubation in a blocking solution containing 1% hydrogen peroxide, 20% normal goat serum, and 1% bovine serum albumin. Sections were then incubated for 48 h in the primary antibodies: Monoclonal mouse anti-parvalbumin (1:5000; P3088; Sigma-Aldrich) and Wisteria floribunda lectin (1:500; L-1516; Sigma-Aldrich). Next, sections were washed in triplicate using TTG, which is a mixture of tris buffered saline, 0.3% Triton-X, and 2% normal goat serum. Sections were next incubated in fluorescent secondary antibodies: AlexaFluor Plus 488 goat anti-mouse IgG (H + L) (1:500; A32723; ThermoFisher Scientific) and Texas Red Streptavidin conjugate (1:1000; S872; ThermoFisher Scientific). After a 2-h incubation, sections were rinsed in duplicate with TTG and in duplicate with TBS, mounted onto a charged microscope slide and coverslipped using EverBrite^TM^ Mounting Medium with DAPI (Biotium, #23002). Negative controls were also included in each stain run to ensure minimal auto-fluorescence and off target secondary antibody binding was maintained. Every stain run included at least one animal from each group, to control for inter-run variability. Slides were stored protected from light at 4 °C until imaging within 5 days.

Prior to any image acquisition, image settings were piloted using samples of adult male rat mPFC and then standardized for all animals. Cross-reactivity between primaries and secondaries was ruled out by applying PNN secondary to sections with only PV primary and vice versa, no cross reactivity was detected—as expected because WFA cannot interact with a non-conjugated secondary antibody such as AlexaFluorPlus 488. Microscope emission capture settings were also adjusted, such that no wavelengths of light were represented in both channels—though Texas Red and AlexaFluorPlus 488 have a very small emission spectra overlap.

Sections containing the mPFC were imaged using an LSM700 confocal microscope at 100x magnification. The z-stack function was used to sample a known z-depth of tissue, resulting in images 320.9 µm x 320.9 µm x 17.66 µm in size. A total of 2,402 images were analyzed. Z-stacks were centered in the center of the z-axis, location of peak signal intensity. Image xy location was standardized, such that all cortical layers—aside from layer I—were represented in all subjects equally. This step is important, as cell density varies across cortical layers. First the mPFC was visualized using DAPI signal, next PV signal was used to locate the division between the prelimbic (PL) and infralimbic (IL) subdivisions of the mPFC. Images were taken from 3 sections, resulting in 12-16 PL images and 4-6 IL images. Images were analyzed in ImageJ using a protocol adapted from Drzewiecki et al. [27]. Briefly, z-stacks were first separated by fluorescent channel, so that PNN and PV intensity could be measured separately. Next, the z-stacks for each channel were projected into a single photo by max intensity (to minimize intensity differences based on signal z-depth). Then PV neurons and PNNs were manually selected as regions of interest (ROIs). Experimenters excluded fluorescent artifacts (denoted with high intensity signal in irregular shapes with little variation in signal throughout) and holes in the tissue (evident with more intense background-WFA-labeling around the hole and complete lack of signal from the center). Experimenters also excluded cells only present in one of the four z-stacks to avoid measuring intensity from a cell whose peak intensity was not captured and to avoid measuring intensity of cells present in other sections. To calculate the intensity of both PV neurons and PNNs, corrections for background fluorescent intensity were made using methods adapted from Slaker et al. [28]. First, the image was processed using the Rolling Ball Radius function, which removes portions of continuous background staining from the entire image. Next, three regions in the background were randomly selected to measure the average intensity and standard deviation of the background staining. The background stain threshold was set as the highest average background intensity measure plus two SDs above this mean. Within each ROI, pixels below the background stain threshold were not used to calculate the mean intensity of PV neurons or PNNs using the “NaN” (Not a Number) function. The stain intensity of each cell in the mPFC was collected and averaged resulting in mean stain intensity for each animal. The number of cells counted in all the pictures was divided by the total volume of the pictures counted from to give PV cell density. PV cell density was then multiplied by the volume of the mPFC to obtain PV cell number, this was then repeated for PNNs. Colocalization was assessed by labeling PV cells based off the presence or absence of a PNN, then diving the total number of PV cells with a PNN by the total number of PV cells. Intensities presented were obtained from split channel images.

### 2.5 Total Neuron Number

Because METH exposure had effects in females, but not in males, Nissl-stained sections were also used to quantify the number of all neurons in females. This was done with an optical disector using StereoInvestigator under a 63x objective and oil immersion, resulting in 630x magnification. The area of the mPFC and its prelimbic and infralimbic subdivisions were traced in two sections (bregma +3.0 to +2.5). Those areas were then randomly sampled with a 75µm x 75 µm x 9µm counting frame, with a top guard zone of 1µm. The number of neurons sampled was then divided by the total volume of the counting frames to achieve neuron density. Neuron density (number of neurons/volume) was then multiplied by the volume of the PL and IL regions of the mPFC, resulting in neuron number in the PL and IL. The number of PV neurons was then divided by the number of total neurons and multiplied by 100 to give the percentage of total neurons positive for PV. Sections used for neuron quantification were of a similar bregma as sections used for PV-PNN quantification (∼ +3.00 to +2.50).

### 2.6 Statistical Analysis

All statistical tests were run using Rstudio, version 4.3.2 (https://www.rstudio.com). Mixed linear effects models were run using the “lmerTest” package for the number of PV neurons and PNNs, as well as for their stain intensity, using two-way ANOVAs for age and treatment with litter as a random factor. Males and females were analyzed separately due to differences in the age of pubertal onset. Significant effects of age were analyzed using Tukey’s post hoc test with a pairwise comparison in the “emmeans” package. Significant interactions between age and treatment were assessed by running one-way ANOVAs in each age group, looking for main effects of treatment with litter as a random factor. Age of pubertal onset was analyzed within the groups that were exposed to METH peri-pubertally (EA females and LA males) using one-way ANOVAs looking for a main effect of treatment, and with litter included as a random factor.

Data from 121 animals was included in the study. A total of 63 males were used in six groups: 10 EA saline, 11 EA METH, 8 LA saline, 11 LA METH, 11 adult saline, and 12 adult meth. There were 58 females in six groups: 10 EA saline, 11 EA METH, 10 LA saline, 7 LA METH, 10 adult saline, 10 adult METH. Two LA METH females were excluded because the entire volume of the mPFC could not be parcellated (final group size n = 7).

### 2.7 Additional Analysis of Intensity

After initial data analysis, intensities of PV cells with a PNN were pooled within female groups to create distributions of PV cell intensity as in Slaker et al., [6]. The data was then plotted using “ggplot2” geom_density in Rstudio. Distributions were compared between treatment groups of the same age using the two-sample Kolmogorov-Smirnov test, ks.test function in Rstudio. Distribution techniques such as this add statistical power with high n, as all cells are pooled within a group. They also increase the chance of discovering low effect size findings, so to combat this, distribution tests were only run to further understand the nature of significant *a priori* comparisons—how METH impacts PV cell intensity in females. In addition, Cohen’s d effect sizes were established for each significant distribution comparison using the cohen’s d function in the “lsr” package in Rstudio.

## 3. Results

### 3.1 Pubertal Onset

Age of pubertal onset was analyzed in groups treated with saline or METH during puberty, early adolescent females and late adolescent males, using one-way ANOVAs split by sex. METH exposure in EA did not alter female pubertal onset, which occurred at P 35.80 ± 2.20 and P 35.70 ± 2.71 for saline and METH exposed animals, respectively. In addition, METH exposure during LA did not alter male pubertal onset, occurring at P 42.00 ± 0.93 and P 42.00 ± 1.48 for saline and METH exposed males, respectively.

### 3.2 PV Cell and PNN Number

#### 3.2.1 PL of the mPFC

The number of PV neurons and PNNs were quantified from the PL (Fig. 2A). In the female PL, a significant age by treatment interaction on the number of PV cells was observed (*F*_2,52_ = 3.33, *p* = 0.045). This interaction was such that EA (*p* = 0.021) and adult females (*p* = 0.022) exposed to METH had more PV cells than their saline exposed counterparts (Fig. 2B) with no significant effects detected in LA females. PNN number was also increased by METH, with a significant main effect of treatment on PNN number in females (*F*_1,52_ = 6.26, *p* = 0.016) (Fig. 2C). The effect of METH on PNN number in the female PL was not significantly modified by age. Age did influence PV number in the female PL (*F*_2,52_ = 4.60, *p* = 0.015), resulting in decreased PV number between EA and adults. A significant main effect of age (*F*_2,52_ = 5.85, *p* = 0.0056) was also detected on the percentage of PV cells surrounded by a PNN, which increased between EA and adult females (*p* = 0.005) (Fig. 2D). In the male PL, there were no main effects of treatment or age nor in their interaction for the number of PV cells, PNNs, and their colocalization (Fig. 2E-G).

**Fig. 2:**
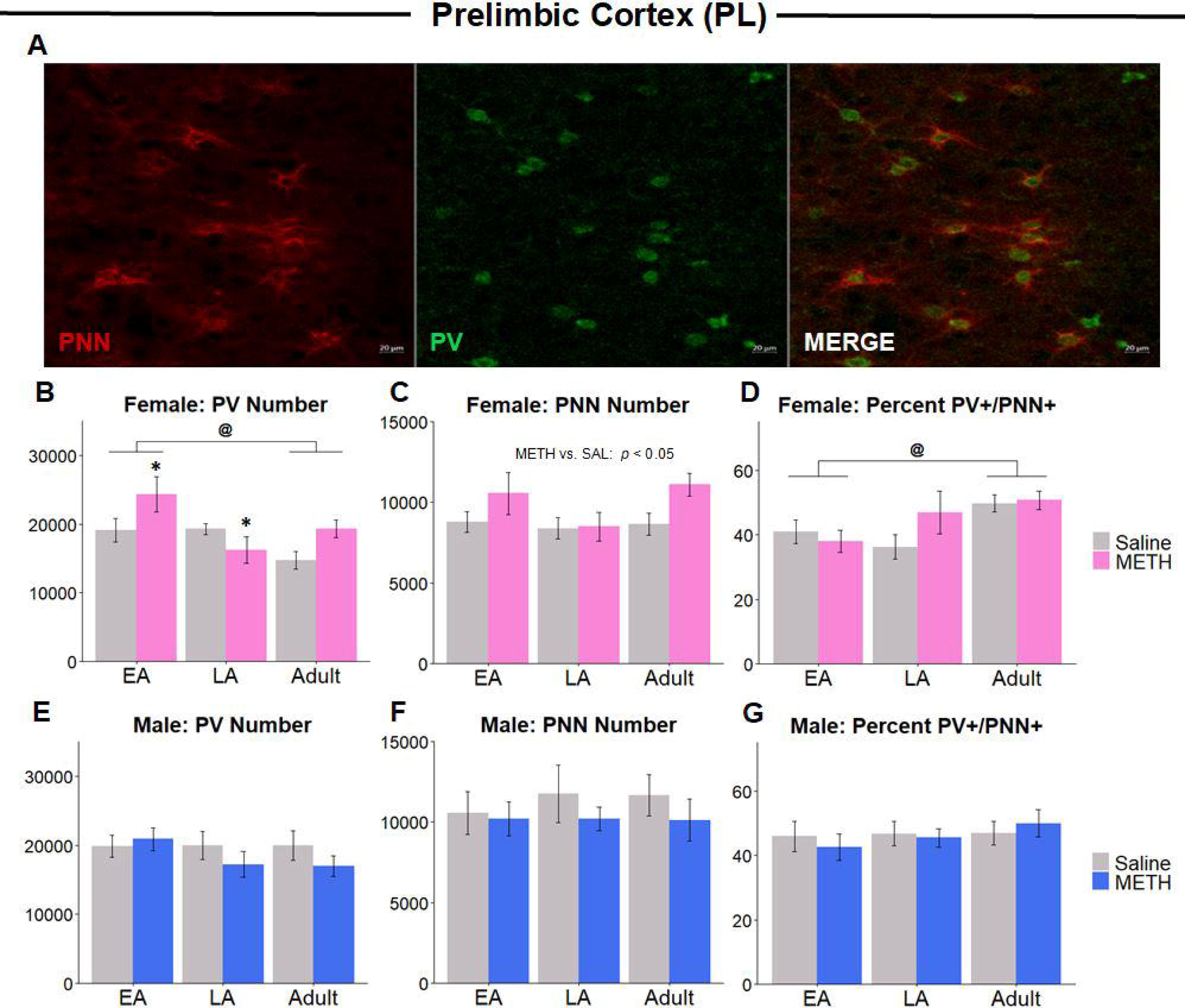
PV+ and PNN number in the PL. (A) Immunofluorescent labeling for PNNs (left), PV (middle), and their colocalization (right). (B) PV cells in the female PL decreased between EA and LA and METH increased the number of PV cells in EA and adult females, but not LA. (C) A significant impact of METH on PNN number was detected in females, with no influence of age. (D) No significant influence of METH on colocalization of PV cells and PNNs was observed, though percent PV enwrapped with a PNN increased between EA and adult females. (E) No influence of METH or age on PV cell number, (F) PNN number, (G) or their colocalization was detected in the male PL. @ = p < 0.05 EA vs. Adult; * = p < 0.05 vs. SAL; EA = early adolescence, LA = late adolescence.

#### 3.2.2 IL of the mPFC

The number of PV cells and PNNs were quantified in the IL (Fig. 3A). In the female IL a significant interaction between age and treatment on PV number was observed (*F*_2, 52_ = 5.45, *p* = 0.0078). The interaction was such that LA METH exposed females had fewer PV cells than saline (*p* = 0.03), while adult METH exposed females had more PV cells than saline (*p* = 0.017). METH exposure also increased the number of PNNs in the female IL, evidenced by a main effect of treatment on PNN number (*F*_1,52_ = 8.72, *p* = 0.005) (Fig. 3C). Like observations in the PL, the effect of METH on PNN number was not modified by age. An age by treatment interaction (*F*_2, 52_ = 4.37, *p* = 0.019) revealed females exposed to METH in LA had a greater percentage of their PV cells enwrapped with a PNN (*p* = 0.04) (Fig. 3D). Age also influenced the number of PV cells within the IL (*F*_2, 52_ = 3.91, *p* = 0.027), with PV cell number decreasing between EA and adult females (*p* = 0.034) (Fig. 3B). Treatment, age and their interaction did not influence the number of PV cells, PNNs, or PV/PNN colocalization in the male IL (Fig. 3E-G).

**Fig. 3:**
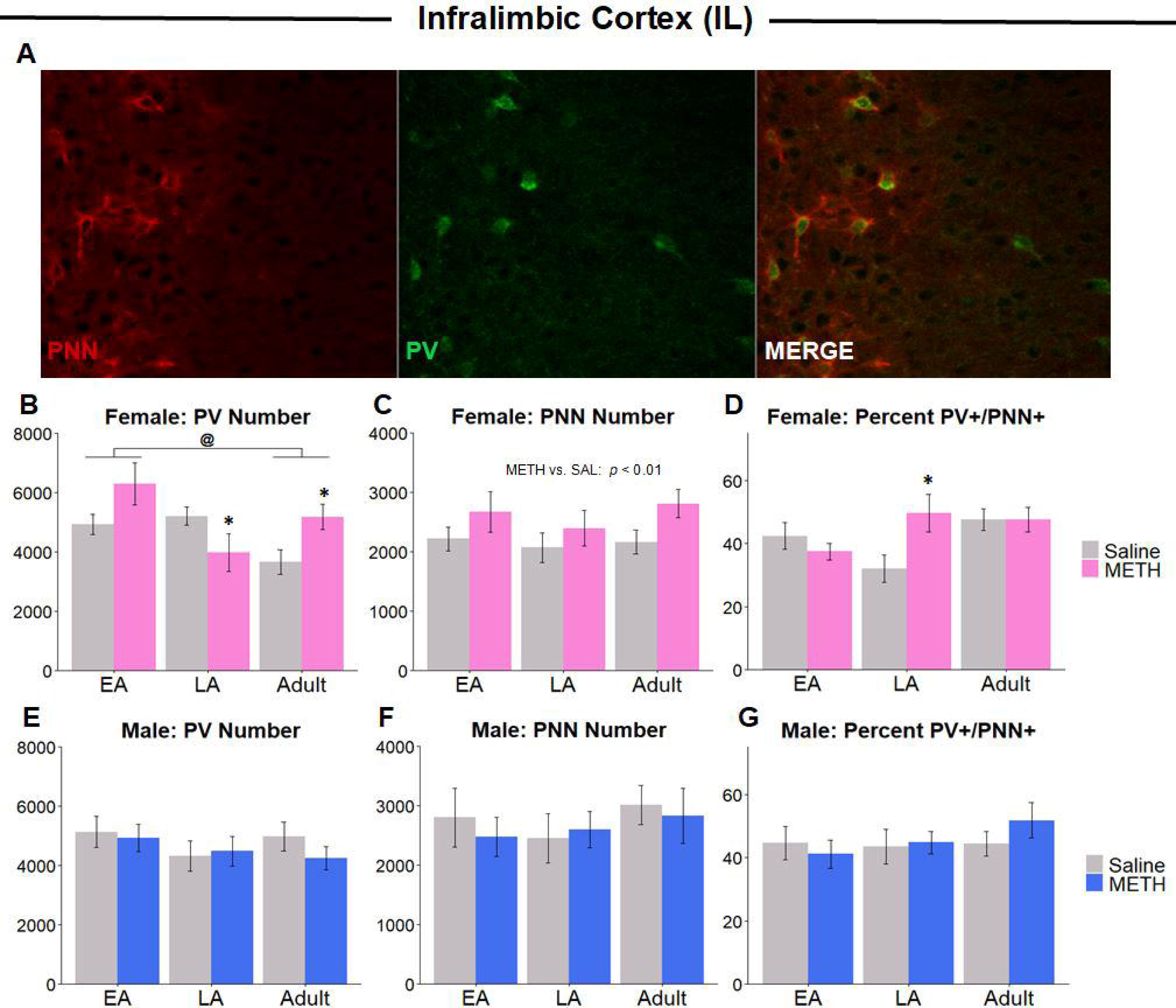
PV+ and PNN number in the IL. (A) Immunofluorescent labeling for PNNs (left), PV (middle), and their colocalization (right). (B) The number of PV cells in the female IL decreased between EA and Adult. METH increased the number of PV cells in adult females but decreased the number of PV cells in LA. (C) A significant impact of METH on PNN number was detected in females, with no influence of age. (D) A significant interaction of METH and age was observed, with LA METH exposed females having a higher percentage of their PV surrounded by a PNN than saline. (E) No influence of METH or age on PV cell number, (F) PNN number, (G) or their colocalization was detected in the male IL. @ = p < 0.05 EA vs. Adult; * = p < 0.05 vs. SAL; EA = early adolescence, LA = late adolescence.

#### 3.2.3 Percentage PV Neurons

Following PV cell number results in females, the number of neurons in the mPFC was quantified to obtain the percentage of neurons positive for PV. No main effects of treatment on percent PV were detected in the PL and IL, however a significant treatment by age interaction was detected in the IL (*F*_2,52_ = 4.35, *p* = 0.019) (Table 1). This was such that adult females exposed to METH had a higher percentage of total neurons positive for PV than saline exposed females (*p* = 0.009).

**Table 1:** Percentage of Total Neurons Positive for Parvalbumin in females (Mean ± SE). #PV Neurons / #Total Neurons; * p < 0.05 vs. saline.

|  |  | *EA* | *LA* | *Adult* |
| --- | --- | --- | --- | --- |
| *PL* | SAL | 3.77% ± 0.24 | 4.01% ± 0.23 | 3.62% ± 0.32 |
| | METH | 4.30% ± 0.31 | 3.59% ± 0.42 | 4.04% ± 0.25 |
| *IL* | SAL | 3.22% ± 0.15 | 3.45% ± 0.25 | 2.86% ± 0.28 |
| | METH | 3.45% ± 0.35 | 2.54% ± 0.35 | 3.34% ± 0.24* |

Abbreviation: PL, prelimbic cortex; IL, infralimbic cortex; EA, early adolescence; LA, late adolescence.

### 3.3 PV Cell and PNN Stain Intensity

#### 3.3.1 PL

In females, METH treatment significantly increased the intensity of PV cells surrounded by a PNN (*F*_1,52_ = 5.79, *p* = 0.021), (Fig. 4A); though intensity of PV cells not surrounded by a PNN was not significantly altered (Fig. 4B). The intensity of PNNs surrounding PV cells increased with age in females (*F*_2,52_ = 13.09, *p* < 0.0001) (Fig. 4C), with the intensity of PNNs on PV cells increasing between EA - LA (*p* = 0.0021) and EA - adult females (*p* < 0.0001). Intensity of PNNs surrounding other cell types was not significantly affected by either treatment or age in the female PL (Fig. 4D).

**Fig. 4:**
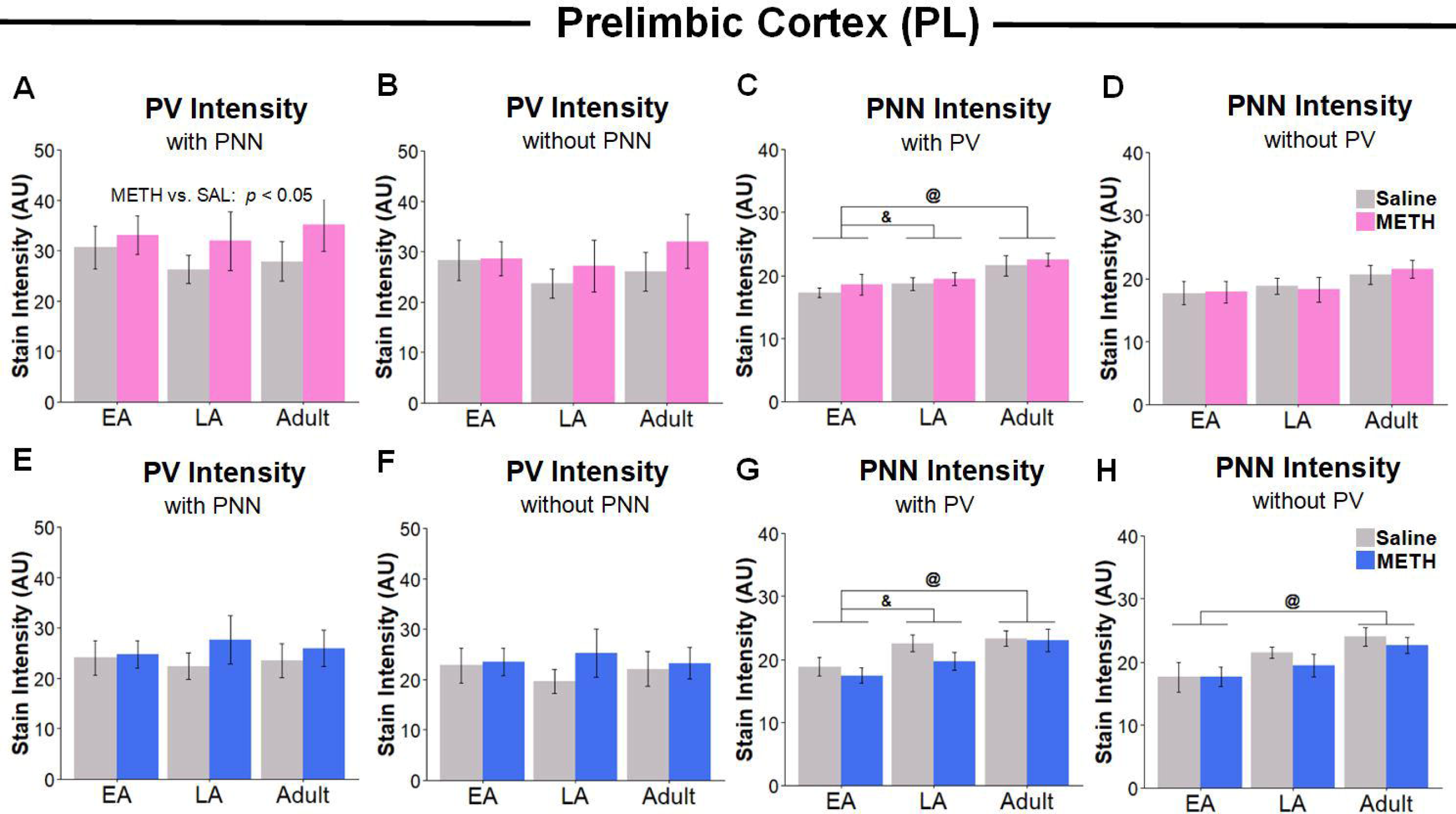
Stain intensity of PV cells and PNNs in the PL. (A & B) In females, METH exposure increased intensity of PV cells surrounded by a PNN but not those without a PNN. (C & D) In females, METH exposure did not alter the intensity of PNNs, either those on PV cells or those on other cell types. (E & F) In males, METH did not alter intensity of PV cells whether surrounded by PNN or not. Intensity of PNNs surrounding PV increased with age. (G & H) In males, METH also did not influence the intensity of PNN staining, though PNN stain intensity increased with age. @ = p < 0.05 EA vs. Adult; & = p < 0.05, EA vs. LA; EA = early adolescence, LA = late adolescence; AU = Arbitrary Units

No main effects of or interactions with treatment on PV cell and PNN intensity were observed in males (Fig. 4E-H). Significant effects of age on the intensity of PNNs surrounding PV cells were noted in males (*F*_2,57_ = 10.21, *p* = 0.00021), with the intensity of PNNs on PV cells increasing between EA - LA (*p* = 0.025) and EA - adult (*p* = 0.0001) (Fig. 4G). In males there was also a main effect of age on the intensity of PNNs on cells not positive for PV (*F*_2,57_ = 7.81, p = 0.0013), with PNN intensity increasing between EA and adult (*p* = 0.008) (Fig. 4H).

#### 3.3.2 IL

In females, a significant main effect of METH on the intensity of PV cells surrounded by a PNN was discovered (*F*_1,52_ = 6.60, *p* = 0.013) (Fig. 5A), this effect was not modified by age. While METH increased the intensity of PV cells surrounded by a PNN in females, PV cells not surrounded by a PNN were not significantly influenced (Fig. 5B). A significant effect of age on the intensity of PNNs surrounding PV cells was noted in females (*F*_2,52_ = 4.65, *p* = 0.015) (Fig. 5C). With the intensity of PNNs on PV cells increasing between EA and adult groups (*p* = 0.017), though PNNs on other cell types were not significantly influenced by age (Fig. 5D).

**Fig. 5:**
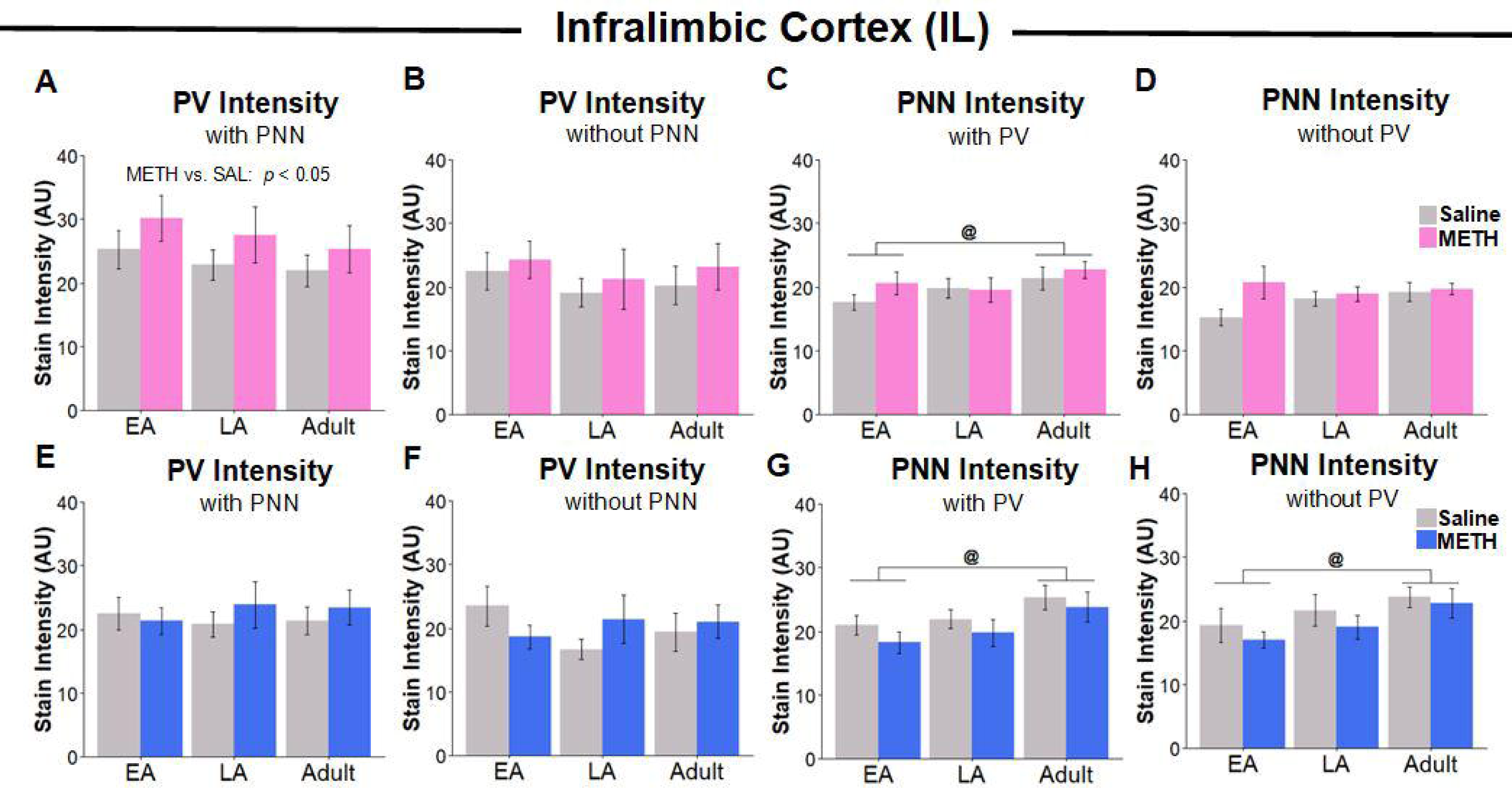
Stain intensity of PV cells and PNNs in the IL. (A & B) In females, METH exposure increased intensity of PV cells surrounded by a PNN but not those without a PNN. (C & D) In females, METH exposure did not alter the intensity of PNNs, either those on PV cells or those on other cell types. Though PNNs surrounding PV cells intensity increased between EA and adult females. (E & F) In males, METH did not alter intensity of PV cells whether surrounded by PNN or not. (G & H) In males, METH also did not influence the intensity of PNN staining, though PNN stain intensity increased between EA and adult. @ = p < 0.05 EA vs. Adult; & = p < 0.05, EA vs. LA; EA = early adolescence, LA = late adolescence. AU = Arbitrary Units

No main effects of or interactions with treatment on PV cell or PNN intensity were observed in males (Fig. 5E-H). A significant effect of age on the intensity of PNNs surrounding PV cells was noted in males (*F*_2,57_ = 5.91, *p* = 0.0051) (Fig. 5G), with the intensity of PNNs on PV cells increasing between EA and adult (*p* = 0.0049). PNNs surrounding other cell types were also influenced by age in the male IL (*F*_2,57_ = 4.52, *p* = 0.016), with the intensity of PNNs on cells not expressing PV increasing between EA and adult groups (*p* = 0.012) (Fig. 5H).

#### 3.3.3 Intensity Distributions

Intensities of PV cells surrounded by a PNN were pooled within groups to create PV cell stain intensity distributions (Fig. 6). In EA females exposed to METH, the distribution of PV cell intensity was significantly altered in the IL (*D* = 0.18, *p* < 0.0001) but not in the PL (*D* = 0.044, *p* = 0.35), suggesting increased number of PV neurons seen in a priori analysis does not change the distribution of PV neuron intensity. The changes noted in early adolescent female PV cell intensity distributions represents a small effect size change (*d* = 0.40). PV intensity distributions were significantly different between METH and saline exposed late adolescent females in both the PL (*D* = 0.17, *p* < 0.0001) and the IL (*D* = 0.25, *p* < 0.0001). These results denote small effect size changes: PL *d* = 0.29 and IL *d* = 0.40. Adult female PV cell distribution was also impacted by METH exposure in both the PL (*D* = 0.15, *p* < 0.0001) and the IL (*D* = 0.12, *p* = 0.014). Again, these results represent small effect size changes: PL *d* = 0.27 and IL *d* = 0.25.

**Fig. 6:**
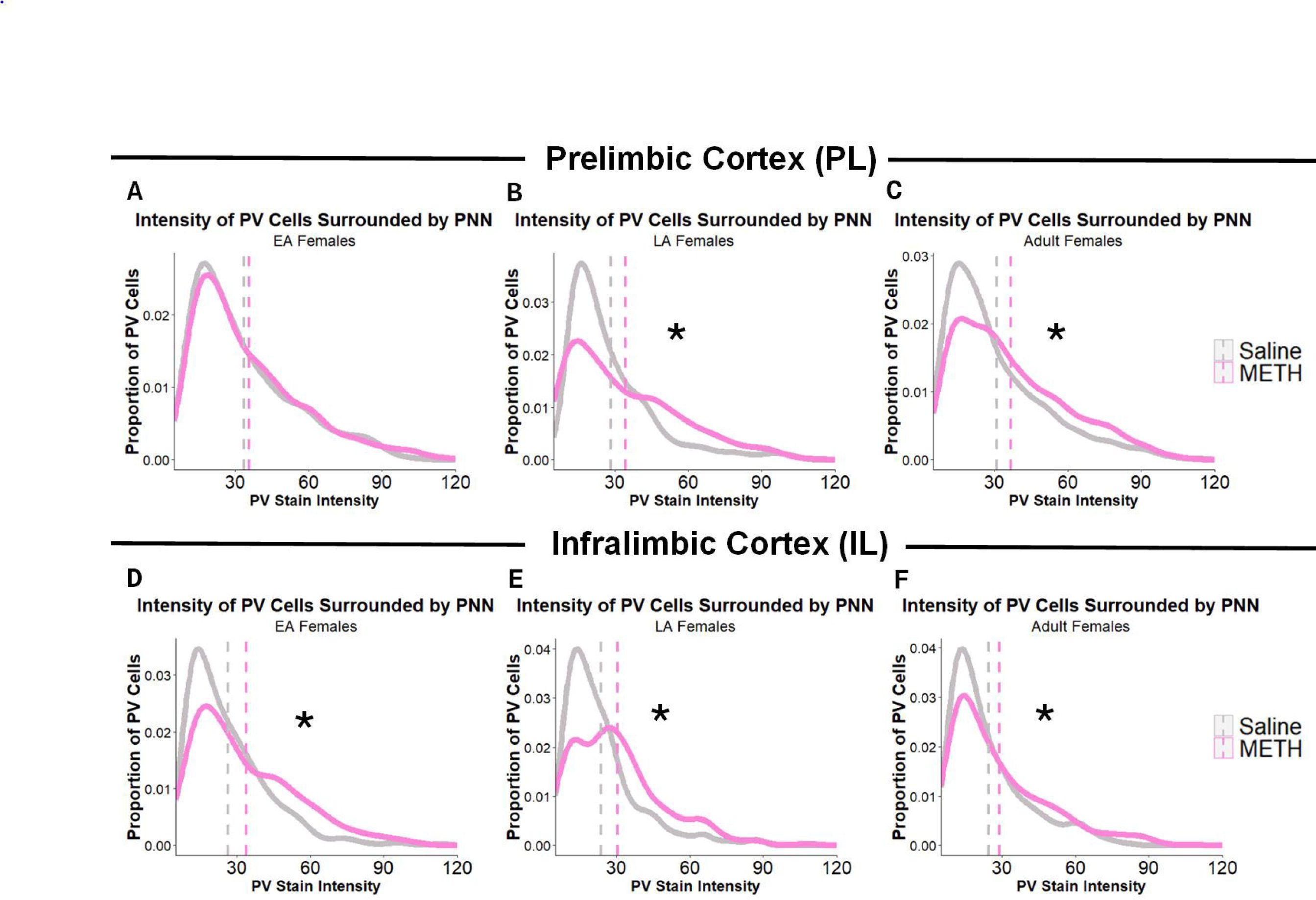
Stain intensity distribution of PV cells in the PL (A-C) and IL (D-F) in females. Early adolescent females (A & D), late adolescent females (B & E) and adult females (C & F). Plotted as proportion of total PV cells(y) of varying intensity (x). * = Saline ≠ METH. Dotted line = group mean.

## 4. Discussion

METH exposure over nine days affected both PV neurons and PNNs in the mPFC during adolescence and adulthood but only in female, not male, rats. Interestingly, the effects of exposure were age-specific such that drug exposure in early adolescence had different effects than exposure in late adolescence. It is also notable that effects of exposure in adult females were more similar to those in early, rather than late, adolescents.

We found that in adulthood, females exposed to METH had more PV neurons than females exposed to saline in both the PL and IL. This result is likely not due to neurogenesis, given the negligible levels during adulthood in the cerebral cortex [29]. Instead, one can speculate that a population of PV-expressing GABAergic neurons has a reduced level of PV expression between the adolescent and adult periods, and this population retains the ability to produce PV in an activity-dependent manner due to METH exposure increasing PV intrinsic excitability and firing rate [5]. This may account for data suggesting that while PV neuron density decreases into adulthood [30,31], GABAergic neuron number does not change [32]. This is further supported by the finding that METH treatment increased the intensity of PV staining, likely bringing the staining of previously sub-detectable PV neurons above the detection threshold. Others have also found increased PV cell expression after an insult in adulthood. In the anterior cingulate cortex, repeated AMPH exposure increased PV density in adult males [33]. PV density also increased after 2 weeks of unpredictable chronic mild stress, specifically in females [34]. This highlights the importance of understanding the nature of protein markers, as intensity of PV stain and therefore PV protein content, directly influences the density of cells quantified. Additionally, loss of a marker may not represent cell death.

Early adolescent females exhibited similar effects of METH as adult females, with increased number of PV neurons in the PL, though the increased number did not reach significance in the IL—likely because of sampling constraints due to IL size. The intensity of PV neurons that were surrounded by PNNs also increased in both PL and IL of early adolescent females exposed to METH. We surmise that females exposed to METH in early adolescence experience a developmental increase in PV expression earlier than those given saline, resulting in a greater number of cells detected. While females exposed to METH in late adolescence had a reversal of this phenotype, with fewer PV cells in the IL. This is the first study to assess the influence of any amphetamine on expression of parvalbumin during adolescence and as a result, at present, we can only speculate on the nature of these effects.

A source of the age-dependent results seen in the present study may be female pubertal onset. Most of the exposure to METH in the early adolescent females was done pre-pubertally, whereas late adolescent females were dosed strictly post-pubertally. While adult females were also exposed post-pubertally, this exposure likely occurred after adolescent cortical reorganization. Therefore, late adolescence may represent a unique period of reorganization and vulnerability of the mPFC. In support of this, work from our laboratory has frequently demonstrated that female pubertal onset influences synaptic pruning, neuronal pruning, and myelination within the rodent mPFC [32,35,36]. Our laboratory has also found that the responsivity of the mPFC to estrogen likely decreases at female puberty which appears to be a catalyst for the reduction of *Esr2* (mRNA for the estrogen receptor β (ERβ)), within the mPFC specifically [27]. In the mPFC, ERβ colocalizes extensively with PV neurons [37] and has been shown to mediate estrogen’s influence on inhibition across the estrous cycle [38]. Given estrogen modulates PV cell activity in adulthood via ERβ, effects of estrogen on PV may be particularly enhanced prior to reduction of *Esr2* in early adolescence. Additionally, following female puberty, GABAergic signaling increases in the mouse cingulate cortex. This increase is blocked by a pre-pubertal ovariectomy, but not by post-pubertal ovariectomy, suggesting reorganization of inhibitory neurotransmission by pubertal onset [39] which is the distinguishing feature of early and late female adolescence in this study.

The different effects of METH exposure in early versus late adolescence indicate that adolescence is not a single entity but rather a series of cellular changes that interact with environmental events such as METH exposure, so that timing is important. The role of timing has also been noted following repeated non-contingent amphetamine exposure, where both males and females treated from P27-45 experienced reduced D1-receptor levels in the mPFC while those exposed from P37-55 did not [40]. Another example of differential timing comes from work on pre-pulse inhibition (PPI), a task that relies on the mPFC [41,42]. PPI is deficient in females that experienced daily restraint stress during early, but not late, adolescence [22]. Interestingly, PPI was also deficient in males when the stress was applied peri-pubertally which coincides with late adolescence in the present study, and males were not affected even later, but post-pubertal, in adolescence. Thus, PPI was deleteriously affected when stress occurred peri-pubertally in both sexes but not after puberty. Others have also seen differential influences based on timing, where 4.0 mg/kg AMPH exposure in female mice from P35-44, but not P22-31, resulted in changes to gene expression within the VTA. Interestingly, in this study males exposed from P22-31, but not P35-44, exhibited similar changes to gene expression within the VTA [43]. This suggests that drug effects can occur at different ages in each sex.

In the present study, females exposed to METH had more PNNs in both the PL and IL; however, this increase seems to be in proportion to the changes in PV neuron number in both the early adolescent and adult exposed groups. Females exposed to METH in late adolescence had a greater percentage of their PV neurons surrounded by a PNN, though no change to the percentage of PNNs on PV neurons. This suggests that the loss of PV expression likely represents PV neurons that did not have a PNN. PNNs have many functions, one of which is giving protection from oxidative stress [44]. PV neurons are prone to oxidative stress due to their activity level [45] and METH has been shown to induce oxidative damage [46,47]. At a speculative level, it is possible that given PNN number temporarily decreases in the mPFC after female puberty [10], PV neurons could be particularly vulnerable to oxidative stress during this time resulting in cell death.

PNN number was not influenced by age in either males or females, contradictory to previous work from our laboratory, where PNN number increased between adolescence and adulthood [10]. Others have also failed to find a significant difference between later adolescent animals and adults [48], suggesting the age-related increase in PNNs may depend mostly on PNN gains prior to P39, the earliest timepoint sampled in the current study. The timing of PNN gains across development may also vary between subjects, given the responsivity to environmental influences displayed by PNNs of the mPFC [49–51]. PNN number did not change with age however PNN intensity increased with age, similar to previous findings [10], suggesting increased accumulation of components of the extracellular matrix within PNNs which is a sign of their maturation [28,52].

PV neuron number and intensity were not influenced by METH in males, suggesting sex specific mechanisms. Studies in adult males have shown modifications to inhibitory neurotransmission during exposure to amphetamines [5,53] and long-term modifications to inhibitory neurotransmission after adolescent stimulant exposure [8,40,54], but females were not examined. These changes to PV neuronal function may not consistently result in changes to the number or density of PV neurons as results in adult males vary [7,55,56].

The current study leads to many questions such as whether the effects within 24 h of METH exposure examined here continue into later stages of life. This is especially important for the increased number of PV neurons detected following exposure during early adolescence since compensation could occur during drug-free late adolescence. Examination of the effects of non-contingent exposure at these ages on later self-administration also should be investigated. Additionally, the role of pubertal onset needs to be explicitly explored. Lastly, the effects of other doses of METH are needed to examine the extent to which the present results generalize.

The present study underscores two important points. One is the importance of using both sexes in drug research as sex differences likely exist at all levels. There are sex differences in self-administration behavior [18,57,58] and long-term cognitive deficits following METH exposure [20]. Second, the present study highlights the importance of not viewing adolescence as a single period, but rather an ongoing set of cellular processes that can interact differentially with drug exposure, making the exact timing of environmental exposures particularly important.

## Statement of Ethics

This study was approved by the IACUC of the University of Illinois protocol number 19113.

## Author Contributions

Amara Brinks: conceptualization, investigation, visualization, writing – original draft; Lauren Carrica: investigation, writing – review and editing; Dominic Tagler: investigation; Joshua Gulley: conceptualization, funding, writing – review and editing; Janice Juraska: conceptualization, funding, supervision, writing – review and editing

## Data Availability Statement

All data generated or analyzed during this study are included in this article. Further enquiries can be directed to the corresponding author.

